# Aligning Brains into a Shared Space Improves their Alignment to Large Language Models

**DOI:** 10.1101/2024.06.04.597448

**Authors:** Arnab Bhattacharjee, Zaid Zada, Haocheng Wang, Bobbi Aubrey, Werner Doyle, Patricia Dugan, Daniel Friedman, Orrin Devinsky, Adeen Flinker, Peter J. Ramadge, Uri Hasson, Ariel Goldstein, Samuel A. Nastase

**Affiliations:** Department of Electrical and Computer Engineering, Princeton University, Princeton, NJ; Department of Psychology and the Neuroscience Institute, Princeton University, Princeton, NJ; Google Research; New York University Grossman School of Medicine, New York, NY; Business School, Data Science Department and Cognitive Department, Hebrew University, Jerusalem, Israel; New York University Tandon School of Engineering, Brooklyn, NY

## Abstract

Recent studies have shown that large language models (LLMs) can accurately predict neural activity measured using electrocorticography (ECoG) during natural language processing. To predict word-by-word neural activity, most prior work has estimated and evaluated encoding models within each electrode and subject—without evaluating how these models generalize across individual brains. In this paper, we analyze neural responses in 8 subjects while they listened to the same 30-minute podcast episode. We use a shared response model (SRM) to estimate a shared information space across subjects. We show that SRM significantly improves LLM-based encoding model performance. We also show that we can use this shared space to denoise the individual brain responses by projecting back into the individualized electrode space, and this process achieves a mean 38% improvement in encoding performance. The strongest improvement was observed for brain areas specialized for language comprehension, specifically in the superior temporal gyrus (STG) and inferior frontal gyrus (IFG). Critically, estimating a shared space allows us to construct encoding models that better generalize across individuals.

## 1 Introduction

Recent advances in the field of natural language processing have showcased the exceptional performance of large language models (LLMs) across various natural language tasks such as text generation, translation, summarization, and question-answering (Devlin et al., 2019; Brown et al., 2020; Manning et al., 2020). In parallel, recent studies in human neuroscience have begun positioning LLMs as computational models of human brain activity during context-rich, real-world language processing (Schrimpf et al., 2021; Caucheteux & King, 2022; Goldstein et al., 2022; Kumar et al., 2022; Toneva et al., 2022). In these works, researchers use encoding models to estimate a linear mapping between internal representations—i.e. embeddings—extracted from an LLM and measurements of human brain activity, word-by-word during natural language comprehension. This simple approach of linearly “aligning” the LLM’s internal feature space to human brain features has yielded remarkably good prediction performance in both functional magnetic resonance imaging (fMRI) and electrocorticography (ECoG). The high spatiotemporal resolution of invasive ECoG recordings, in particular, promises to provide finer-grained insights into shared representations and processes between LLMs and the brain (Goldstein et al., 2022; Cai et al., 2023; Goldstein, Wang, et al., 2023; Mischler et al., 2024; Zada et al., 2023; Goldstein et al., 2024).

When exposed to the same natural language stimulus, such as a spoken story, human neural activity converges on stimulus features ranging from basic acoustic attributes to more complex linguistic and narrative elements (Lerner et al., 2011; Honey et al., 2012; Hasson et al., 2015). However, while a coarse alignment exists across individual brains (Nastase et al., 2019; 2021), the finer cortical topographies for language representation exhibit significant idiosyncrasies among individuals (Fedorenko et al., 2010; Nieto-Castañón & Fedorenko, 2012; Braga et al., 2020; Lipkin et al., 2022). To address this, hyperalignment techniques have been developed in fMRI research to aggregate information across subjects into a unified information space while overcoming the misalignment of functional topographies across subjects (Haxby et al., 2011; Chen et al., 2015; Guntupalli et al., 2016; Haxby et al., 2020; Feilong et al., 2023). Unlike fMRI, where there is putative voxelwise correspondence during acquisition, ECoG presents a more difficult correspondence problem because each subject has a different number of electrodes in different locations (with placement guided by clinical considerations, not research goals). Thus, how to best aggregate electrodes across individuals is a matter of ongoing research (e.g., Owen et al., 2020). For this reason, encoding models are typically constructed separately at each electrode within individual subjects and are not assessed for their generalization to new subjects (e.g., Schrimpf et al., 2021; Goldstein et al., 2022).

In this paper, we measured the neural responses of eight ECoG subjects implanted with invasive intracranial electrodes while they listened to a natural language stimulus. We develop a shared response model (SRM; Chen et al., 2015) to aggregate neural activity and isolate a stimulus-driven shared feature space that is shared across individuals. In parallel, we use LLMs to extract contextual embeddings for each word of the podcast. We then build encoding models to estimate a linear mapping from the contextual embeddings to the shared neural features (Van Uden et al., 2018; Nastase et al., 2020). We show that the SRM yields significantly higher encoding performance than the original individual-specific electrodes. Moreover, we show that we can use this shared space to “denoise” individual subject responses by projecting from the shared space back into the individual electrode space. We find that the SRM-reconstructed data yields the largest improvement in brain areas specialized for language comprehension. Finally, we demonstrate that the SRM allows us to construct encoding models that better generalize across subjects.

## 2 Materials AND Method

### 2.1 Data Collection AND Processing

We recorded the neural activity of eight participants (4 reported female, 20–48 years) using ECoG. Participants were presented with a 30-minute audio podcast “So a Monkey and a Horse Walk Into a Bar, Act One: Monkey in the Middle” from the *This American Life* podcast. We manually transcribed the story and aligned it to the audio by labeling the onset and offset of each word. An independent listener manually evaluated the alignment. There were a total of 5,013 words in the podcast. Using the Hugging Face environment (Wolf et al., 2019), we supplied the transcript to the large language model GPT-2 XL (Radford et al., 2019). For each word, a 1600-dimensional contextual embedding was extracted from the final layer of the model. The meaning of the embedding for each word (excluding the first word) was contextualized by the preceding words in the podcast stimulus. These embeddings were reduced to 50 dimensions using principal component analysis (PCA) for the core encoding analyses, based on Goldstein et al. (2022).

For ECoG data collection, 917 electrodes were placed on the left hemisphere and 233 on the right hemisphere. The ECoG data were sampled at 512 Hz. Line noise harmonics were excluded. We used a band-pass filter to extract activity in the high gamma range of 70-200 Hz.

### 2.2 Encoding Models

We use contextual word embeddings to predict held-out neural data for individual electrodes or SRM features (see below). First, we extracted gamma power in 200 ms windows at 161 lags ranging from -2000 ms to +2000 ms in 25 ms increments for epochs indexed to each word’s onset. For each lag at a given electrode, we then estimated electrode-wise encoding models using ordinary least-squares multiple linear regression: this yields a linear mapping to predict word-by-word neural activity from the associated contextual embeddings (Naselaris et al., 2011; Huth et al., 2016). We employ 10-fold cross-validation to assess the performance of these models in predicting neural responses for held-out, temporally-contiguous segments of the stimulus. We evaluated out-of-sample prediction by computing the Pearson correlation between the predicted and the actual signal for each held-out test set.

In comparing the encoding performance alignment methods, we performed paired t-tests between the two correlation scores across folds for each lag. To correct for multiple tests across 161 lags, we control the false discovery rate (FDR) at .01 (Benjamini & Hochberg, 1995). Lags with FDR less than .01 are considered to be significant.

### 2.3 Electrode Selection

To select a subset electrodes involved in language processing, following Goldstein et al. (2022), we first estimated encoding performance using non-contextual GloVe embeddings (Pennington et al., 2014) at 161 lags ranging from -2000 ms to +2000 ms in 25 ms increments for epochs indexed to each word’s onset. To evaluate the statistical significance of GloVe-based encoding performance, we performed a randomization test. For each of the electrodes and lags, we randomized the phase of the signal so as to disrupt the temporal alignment while preserving the autocorrelation, then re-estimated the GloVe based encoding models. We repeated this procedure for 5,000 phase randomizations to construct a null distribution from the maximum encoding performance across lags for each electrode. We calculated p-values for each electrode as the percentile of the actual encoding performance relative to 5,000 phase-randomized samples from the null distribution. We controlled the false discovery rate (FDR; Benjamini & Hochberg, 1995) at *q* = .01 across electrodes. If the *q*-value of the electrode was less than .01, it was selected for further analysis. This yielded 184 electrodes (150 in the left hemisphere, 34 in the right hemisphere) across subjects (see Table S1 for a detailed description of electrode coverage).

### 2.4 Shared Response Model

Although all the subjects listened to the same story, both the placement of their electrodes and the functional properties of similarly placed electrodes will tend to differ from individual to individual. We use a shared response model (SRM; Chen et al., 2015) to aggregate ECoG data across subjects into a common information space that accounts for different electrode placement and functional topographies across individuals. SRM learns subject-specific transformations that map from each subject’s idiosyncratic functional space into a shared space based on a subset of training data, then uses these learned transformations to map a subset of test data into the shared space.

To clarify this, let 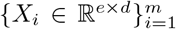 be the training data (e electrodes over d time points) for m subjects. We use this training dataset to learn subject-specific bases *W*_*i*_ ∈ *ℝ*^*e×k*^ (where *k* is a hyperparameter that corresponds to the number of components in the new, shared space) and a shared matrix *S* ∈ *ℝ*^*k×d*^, such that *X*_*i*_ = *W*_*i*_*S* + *E*_*i*_, where *E*_*i*_ is an error term corresponding to deviation from the subject’s original brain activity. The bases *W*_*i*_ represent the individual functional topographies, while S represents latent features that capture components of the response that are shared across subjects. For the solution to be unique, *W*_*i*_ is subject to the constraint of linearly independent columns and *W*_*i*_ is assumed to have orthonormal columns, 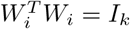 (Chen et al., 2015). The following optimization problem is solved to estimate *W*_*i*_ and the shared response *S*:

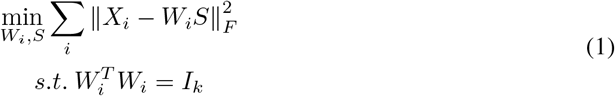

The *S* and *W* parameters of the SRM model are jointly estimated using a constrained EM algorithm. We can utilize the learned subject-specific bases to project data from shared space back into the individual shared response subspace (*S*_*i*_) to reconstruct a “denoise” version of the data in the original electrode space *X*_*i*_:

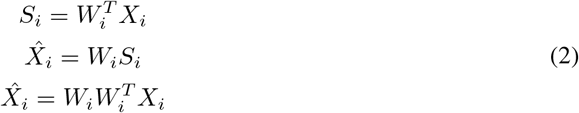

## 3 Results

### 3.1 Linguistic Encoding IN Shared Space

We estimated a shared response model (SRM; Chen et al., 2015) on a training subset of the data (9 out of 10 segments of the story stimulus) across 8 subjects with 5 shared features (hyperparameter *k* = 5). We selected *k* = 5 shared features to maximize the number of eligible subjects, given that subject S4 had only eight electrodes survive the electrode selection procedure. This fitting procedure yields a shared space and the corresponding subject-specific weights (*W*_*i*_). We projected each subject’s training data into the shared space and averaged the reduced-dimension data across subjects (*S*_*train*_). Next, we estimated encoding models using the same training data, comprising the average word-by-word time series across subjects in the reduced-dimension shared space. We used linear least-squares regression to estimate a weight matrix to predict the shared neural activity from contextual embeddings (reduced to 50 dimensions using PCA) extracted from GPT-2 XL (Radford et al., 2019). To evaluate encoding model performance, we first use the subject-specific weights *W*_*i*_ to project the test data (corresponding to the left-out test segment of story stimulus) into the shared space estimated from the training data, and average across subjects: 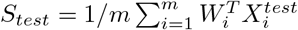

We then use the encoding weights estimated from the training set to generate model-based predictions of neural activity in shared space from the contextual embeddings for the left-out training segment of the story stimulus. We evaluate these model-based predictions by computing the Pearson correlation between predicted and actual neural activity for each shared feature. In this way, both the shared response model and the encoding models are estimated and evaluated within the same 10-fold cross-validation procedures (Van Uden et al., 2018; Nastase et al., 2020). We repeated this analysis for lags from 2000 ms before word onset to 2000 ms after with a 25 ms stride, fitting and evaluating separate encoding models at each lag.

When using contextual embeddings to predict shared features, we observed strong encoding performance with peak accuracy (averaged across shared features) of 0.38 roughly 200ms after word articulation (Fig. 1A). SRM dramatically outperforms typical electrode-wise encoding performance using the same embeddings and cross-validation scheme (Fig. 1B). This improvement, however, could be driven by the fact SRM reduces dimensionality by aggregating signals across electrodes. As a control analysis, we instead aggregated electrodes across subjects using principal component analysis (PCA) with dimensionality *k* = 5 and reassessed encoding performance in the PCA-based shared space. PCA similarly reduces dimensionality with the same orthogonality constraint as SRM, without aligning individual subjects into a shared feature space. We found that SRM achieves significantly higher encoding performance than PCA (*p* < .01, FDR corrected; Fig. 1A). This control analysis shows that the stronger encoding performance is not simply due to the decreased dimensionality of the shared space.

**Figure 1:**
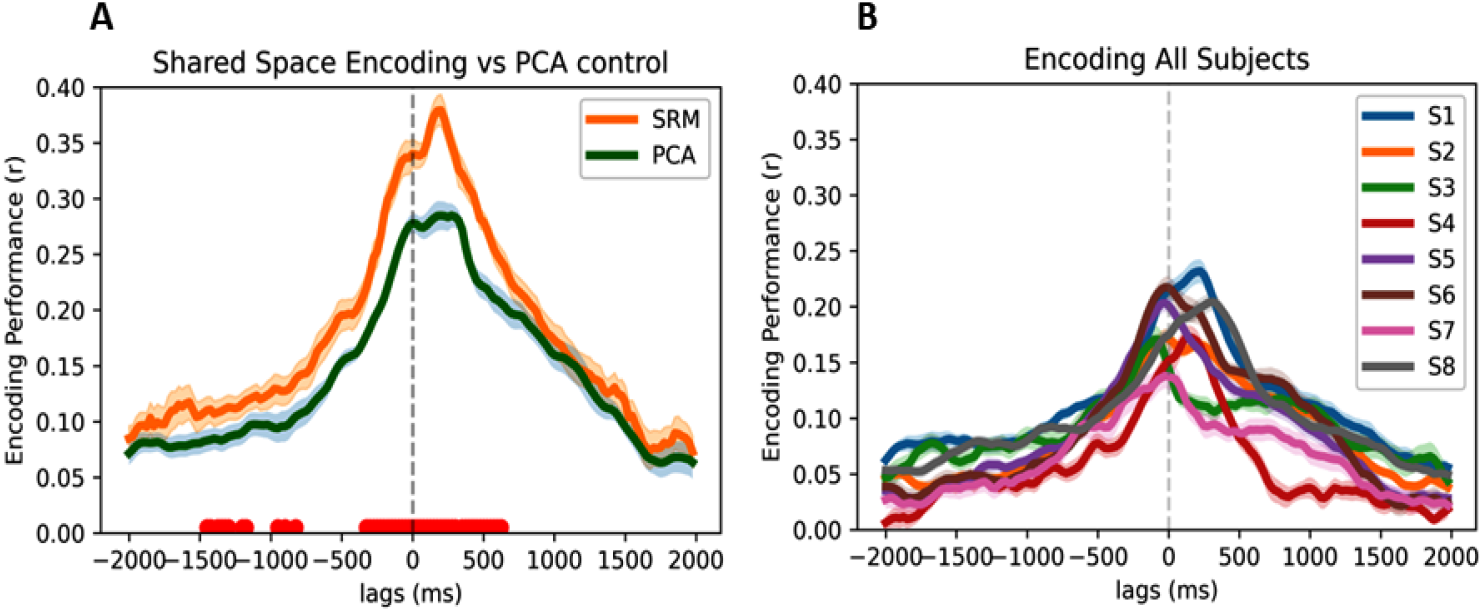
SRM improves model-based encoding performance. (**A**) Encoding model performance based on SRM (orange) and control analysis based on PCA (blue) with matched dimensionality (*k* = 5). As a control analysis, PCA aggregates neural signals across subjects with the same dimensionality reduction and the same orthogonality constraint, but does not align neural response trajectories across subjects. Encoding performance is averaged across features. The red dots at bottom indicate lags with a significant difference between SRM and PCA-based encoding model performance across folds (FDR controlled at .01). Error bands indicate standard error of the mean across cross-validation folds. (**B**) Encoding model performance based on the original neural activity in each subject (*N* = 8). Encoding performance is averaged across electrodes within each subject.

### 3.2 Reconstructing Electrode Activity VIA THE Shared Space

We hypothesize that projecting an individual subject’s neural activity into the reduced-dimension shared space and then back into electrode space will effectively denoise the individual-subject data and increase encoding model performance. First, we transform the individual subject data into the reduced-dimension shared subspace *S*_*i*_ by multiplying it with the learned, subject-specific weights from SRM training. We then use the transpose of the subject-specific weights to reconstruct their electrode data for both the training and the test sets, as shown in equation 2. Then, we perform an encoding analysis for each subject using the SRM-reconstructed data and compare it with the encoding performances using the subjects’ original neural data (Fig. 2). The SRM-reconstructed data significantly improved encoding performance at numerous lags for each subject (*p* < .01 for all subjects, FDR corrected), with an average 38% improvement in peak model performance across subjects.

**Figure 2:**
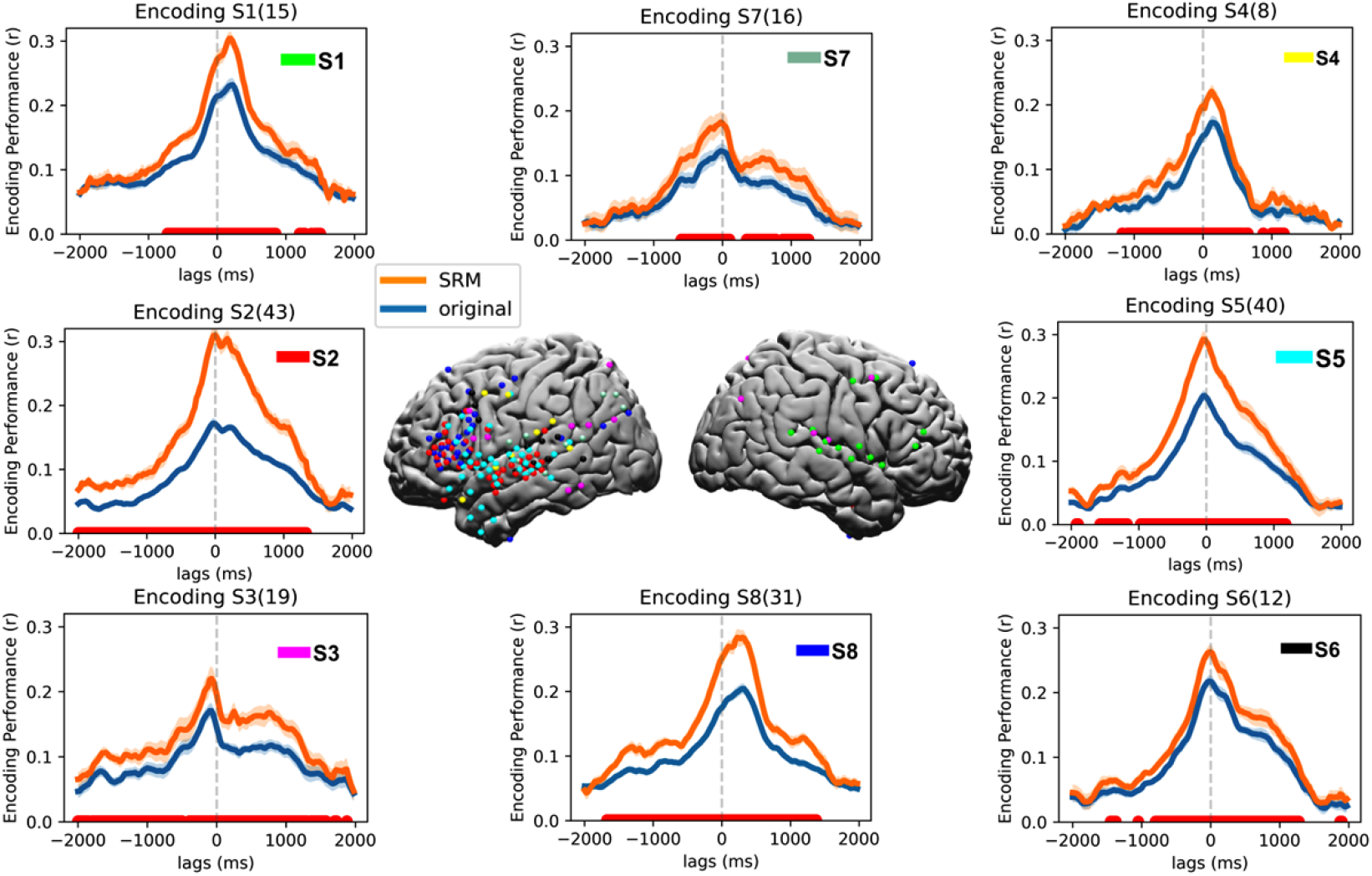
Reconstructing electrode activity via the shared space. At center, electrode placement is shown for all subjects (*N* = 8). Electrode-wise encoding performance is shown for each subject-based electrode activity reconstructed from the shared space (orange) and original electrode activity (blue). Encoding performance is averaged across electrodes within each subject. Error bands indicate standard error of the mean encoding performance across folds. The red markers at bottom indicate lags with a significant difference between encoding performance for SRM-reconstructed and original electrodes across folds (FDR controlled at .01)

### 3.3 Localizing Improved Encoding Performance WITH SRM Reconstruction

To map out which brain regions improve most when reconstructing electrode activity from the shared space, we quantified the difference in encoding model performance between SRM-reconstructed data and the original data for each electrode separately (Fig. S1). Qualitatively, the largest improvements were found in the inferior frontal gyrus (IFG) and the superior temporal gyrus (STG). Table 1 reports the number of electrodes at varying ranges of improvement. Out of 184 electrodes, encoding performance nominally improved in 168 electrodes when reconstructed from the shared space, with a maximum improvement of 0.33. We further examined improvements in encoding performance of SRM-reconstructed data for different areas of the language network (Fig. 3). We observed that SRM reconstruction yields significantly better encoding performance compared to the original electrode data in IFG, anterior STG (aSTG), and middle STG (mSTG) (*p* < .01, FDR corrected). While caudal STG (cSTG), angular gyrus (AG), and temporal pole (TP) show nominal improvement in encoding performance, these improvements are not significant after correcting for multiple lags; this may in part be due to the relatively fewer number of electrodes in these areas.

**Table 1:** Improvement in electrode-wise encoding performance with SRM-reconstructed versus original electrode data. Differences were computed between the respective maximum encoding performance values across lags for SRM-reconstructed and original electrode data.

| $r_{SRM} - r_{original}$ | Number of electrodes |
| --- | --- |
| > 0.20 | 24 |
| 0.15–0.20 | 12 |
| 0.10–0.15 | 43 |
| 0.05–0.10 | 50 |
| 0.01–0.05 | 39 |

**Figure 3:**
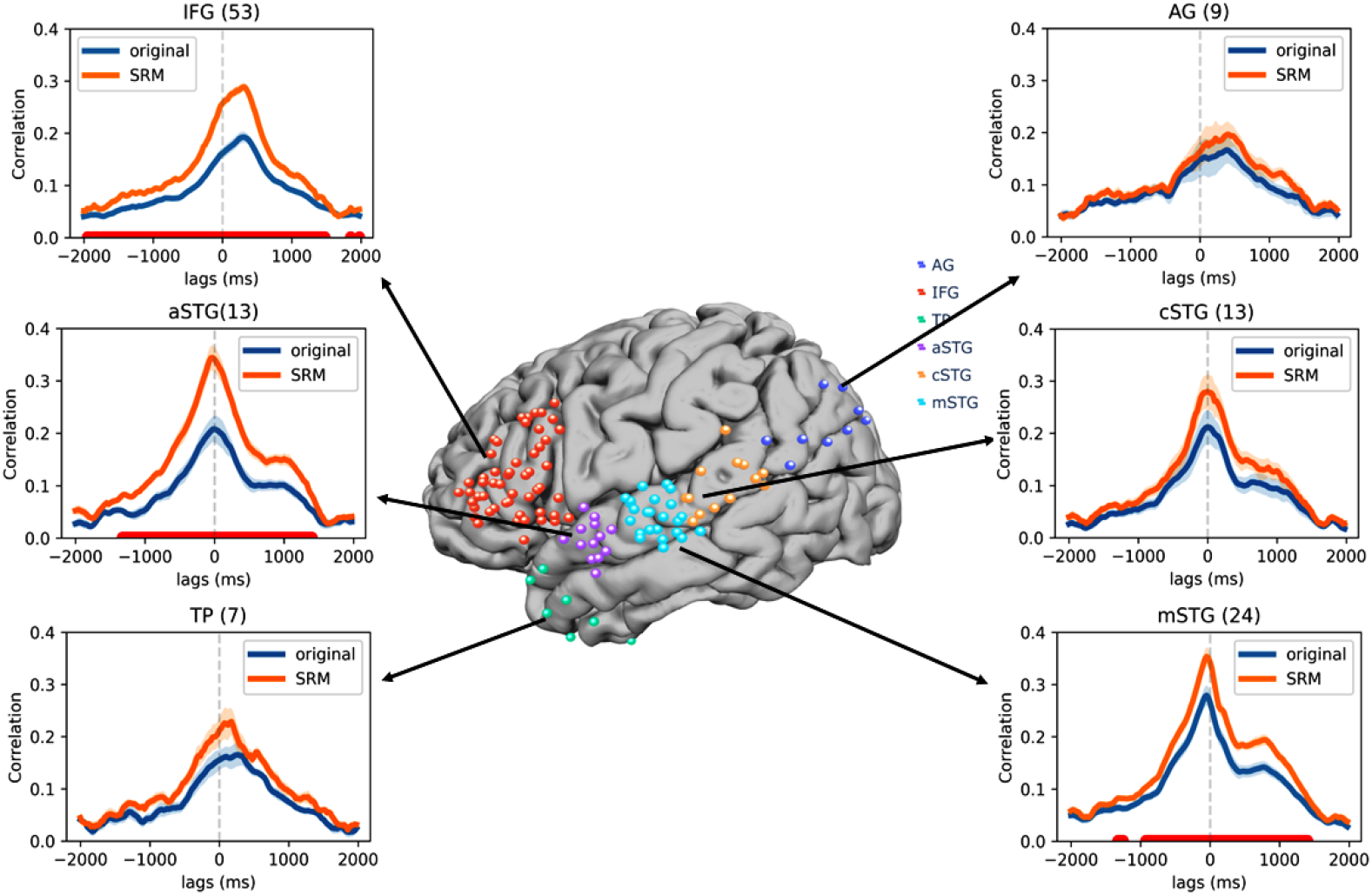
Comparison of encoding performance for SRM-reconstructed data and original electrode data for different regions of the language network. At center, electrode placement is shown for all subjects (*N* = 8). Electrode-wise encoding performance values for lags spanning -2000 ms to +2000 ms lags are shown for each brain area based on electrode activity reconstructed from the shared space (orange) and original electrode activity (blue). Encoding performance is averaged across electrodes within each brain area. Error bands indicate standard error of the mean encoding performance across folds. The red markers at bottom indicate lags with a significant difference between encoding performance for SRM-reconstructed and original electrodes across folds (FDR controlled at .01 across lags). cSTG: caudal superior temporal gyrus, mSTG: middle superior temporal gyrus, aSTG: anterior superior temporal gyrus, TP: temporal pole, AG: angular gyrus, IFG: inferior frontal gyrus.

### 3.4 Generalizing Encoding Models ACROSS Subjects VIA THE Shared Space

In the previous analyses, we showed that SRM can improve encoding performance—but, like in prior work (e.g. Goldstein et al., 2022), those encoding models were estimated and evaluated in individual subjects. The shared space captures the shared, stimulus-driven features of brain activity and retains subject-specific mappings to and from the shared space. SRM should therefore allow us to build encoding models that generalize to new subjects who have received the same stimulus (Van Uden et al., 2018; Nastase et al., 2020). To test this hypothesis, we estimated both SRM and encoding models in a subset of training subjects (for a training segment of the story stimulus), then evaluated encoding model performance on a left-out subject (for the left-out test segment of the story). We first estimate a shared space (*S*) for *N* 1 training subjects based on the training segments of the story. In this case, SRM training data does not include the neural data of the test subject. We can estimate encoding models from the *N* 1 training subjects in this shared space.

Next, we must estimate a transformation to project the test subject’s data into the shared space derived from the training subjects. The shared space (*S*) derived from the training subjects is used as a template, and we calculate a subject-specific weight matrix *W*_*j*_ to rotate the left-out subject *j* into the pre-existing shared space, using the data 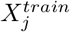 from the training segments of the story. To achieve this, we minimize the mean squared error 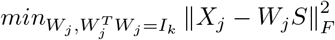 to find *W*_*j*_. The shared space *S* is not affected by aligning a left-out subject in this way. Now, we transform the test subject’s neural activity for the test segment of the story into the shared space (estimated from other subjects) using *W*_*j*_ estimated from the training segments of the story: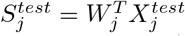. Finally, we evaluate the encoding models estimated from other subjects’ data and training segments of the story. We use the encoding weights trained on the shared space (*S*), combined with the embeddings for the test segment to generate predictions for the left-out subject 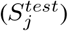. We carry out this process for each lag for all subjects (leave-one-subject-out).

Using SRM, we obtain cross-subject encoding performance (Fig. 4a, orange) comparable to the performance observed when encoding models are estimated and evaluated in individual subjects (Fig. 1b). For a more direct comparison, we implemented a control analysis using PCA: we estimate PCA across *N −* 1 training subjects to learn a PCA-based reduced-dimension space (with matching dimensionality and orthogonality constraints to SRM) from the training story segments; we then calculate a *W* transformation like above to project the test subject onto reduced-dimension PCA reduced space using the left-out subject’s training story segments. Finally, we estimate encoding models in the reduced-dimension PCA space from the training subjects and the training story segments. We project the left-out subject’s test segment into the shared space to evaluate the model-based predictions. Cross-subject encoding performance is nominally better with SRM than with PCA.

**Figure 4:**
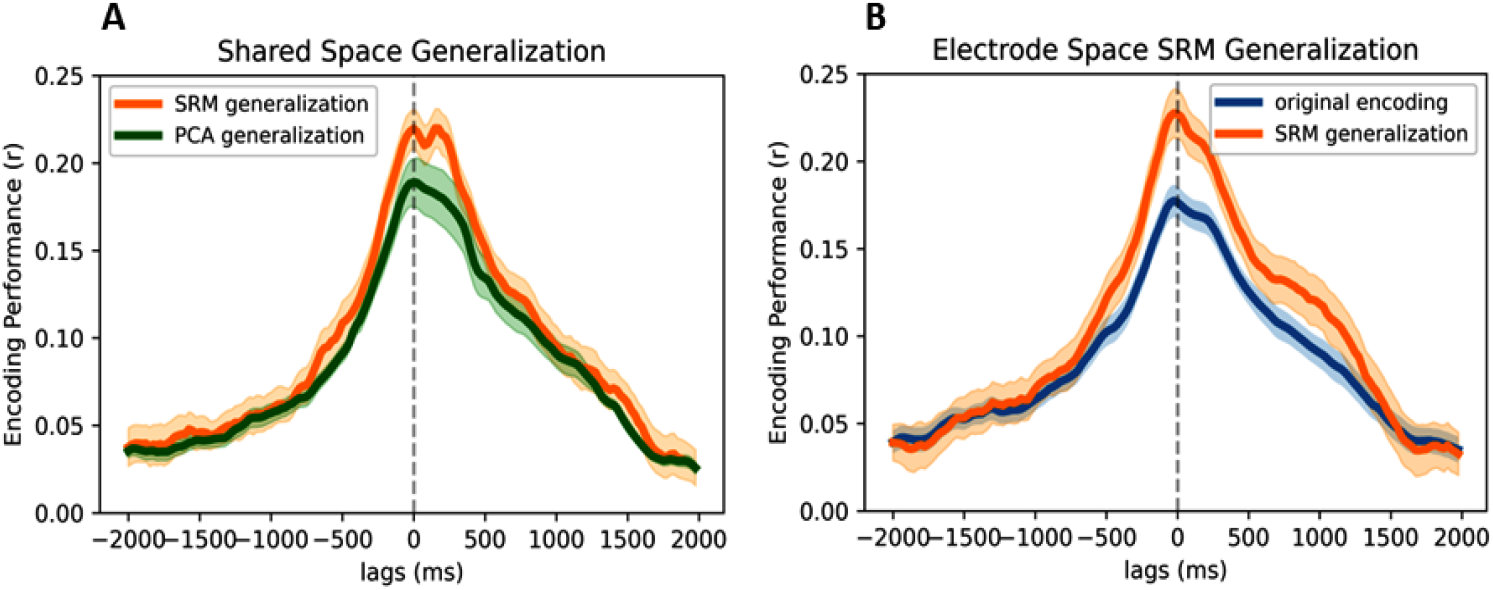
Cross-subject encoding performance via the shared space. In cross-subject encoding, both SRM and encoding models are estimated from *N −* 1 subjects and model-based predictions are tested against a left-out subject. (**A**) Cross-subject encoding performance in SRM-based shared space (orange) compared to PCA control. Encoding performance is averaged across features in shared space. (**B**) Cross-subject encoding performance in the test-subject’s SRM-reconstructed electrode space compared to within-subject encoding performance in original electrode space. Encoding performance is averaged across subjects and SRM features (a) or electrodes (b). Error bands indicate standard error of the mean encoding performance across subjects.

To extend this cross-subject encoding analysis from the reduced-dimension shared space to the original electrode space, we first project *N −* 1 subjects to a SRM shared space (*S*) using the training segments of the story. Next, we calculate the weight matrix *W*_*j*_ to rotate the left-out subject *j* into the shared space. Now we can use *W*_*j*_ to project data from *N −* 1 from the shared space back into the test subject’s space: *X*^*train*^ = *W*_*j*_*S*. This allows us to estimate encoding models based strictly on other subjects’ data in the test subject’s original electrode space: 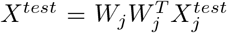. Cross-subject encoding performance in across all the test subject’s SRM-reconstructed electrode space nominally outperforms within-subject encoding models in the original electrode space (Fig. 4b). In both of these analyses, we show that an SRM estimated from *N −* 1 subjects can be used to find a set of shared features that generalize to a new subject with a different number and placement of electrodes. Given a shared stimulus, SRM can provide a robust enough linkage across disparate, individual-specific electrodes to allow us to build encoding models that generalize to a left-out individual.

### 3.5 Quantifying Shared Information ACROSS Subjects

How well can we reconstruct a novel subject’s neural responses to a novel stimulus based on the neural activity of other subjects? To quantify the quality of the shared space without reference to an encoding model, we estimated a shared space based on the training segments of the story in *N –* 1 subjects, then reconstructed neural activity for the left-out test segment in a left-out subject. We then correlate the reconstructed neural activity for the test subject *j* with the subject’s actual neural activity. High correlations indicate that the shared space robustly captures shared information that generalizes across subjects. To elaborate, first, we train an SRM model on the training data for *N −* 1 subjects except *j* using Eq. 1. Then, using the training data *X*_*j*_ for subject *j*, we find the matrix *W*_*j*_ mapping subject *j* into the pre-existing shared space by minimizing the mean squared error of 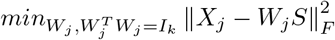 .

Next, we average the shared responses for the test segment across *N −* 1 subjects except *j* using 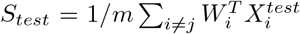 . With this shared response for the test data, we reconstruct the test data for subject *j* (based strictly on data from *N −* 1) by 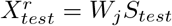. Finally, we calculate the correlation across words between 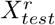 And 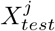 for each electrode. We repeat this process for each test subject for all the test segments at the word onset. We find that SRM-based reconstruction based on the neural activity of other subjects yields .25 correlation on average (Table 2).

**Table 2:** SRM reconstruction quality based on other subjects’ data transformed via the shared space into the test subject’s electrode space. Correlation between SRM-reconstructed and original test data (with standard error of the mean correlation across test sets).

| Subject | Correlation between SRM-reconstructed and original data |
| --- | --- |
| S1 | $0.28 \pm 0.008$ |
| S2 | $0.21 \pm 0.006$ |
| S3 | $0.21 \pm 0.012$ |
| S4 | $0.23 \pm 0.011$ |
| S5 | $0.29 \pm 0.008$ |
| S6 | $0.32 \pm 0.009$ |
| S7 | $0.20 \pm 0.009$ |
| S8 | $0.23 \pm 0.007$ |
| Average | $0.25 \pm 0.009$ |

### 3.6 Exploring SRM Encoding ACROSS DIfferent Models Parameters

Lastly, we explored encoding performance in the shared space across several different sets of model features. We first examined how encoding model performance varies across layers for GPT-2 XL:
we extracted contextual embeddings from all 48 layers of GPT-2 XL and repeated our encoding analysis for both shared features and original electrodes at lags ranging from –2000 ms to +2000 ms relative to word onset. In both cases, we found that intermediate layers yield the highest prediction performance in human brain activity (Fig. S2a,S2b), consistent with prior work (Schrimpf et al., 2021; Caucheteux & King, 2022; Goldstein, Ham, et al., 2023; Kumar et al., 2022). Next, we evaluated encoding models for two different types of word embeddings: contextual (GPT-2 XL) and non-contextual (GloVe) embeddings (Pennington et al. 2014; note that GloVe encoding was initially used to select electrodes). We found that contextual embeddings yield dramatically higher encoding performance that non-contextual embeddings, both for original electrode data and in the shared space (Fig. S2c,S2d; similarly to prior work, e.g., Schrimpf et al. 2021; Goldstein et al. 2022; Kumar et al. 2022. Finally, we repeat our SRM encoding analysis with several open source GPT models ranging from 125M to 20B parameters: neo-125M, large-774M, neo-1.3B, XL-1.5B, neo-2.7B, neo-20B. SRM yields improved encoding performance for all models and we observe a weak trend consistent with previously reported results (Antonello et al., 2023) where larger models yield better encoding performance (Fig. S2e, S2f).

## 4 Discussion

Many recent studies have begun to employ encoding models to predict neural responses during natural language processing using contextual embeddings derived from LLMs (Schrimpf et al., 2021; Caucheteux & King, 2022; Goldstein et al., 2022; Toneva et al., 2022; Cai et al., 2023; Goldstein, Wang, et al., 2023; Mischler et al., 2024; Zada et al., 2023). Our study demonstrates that aligning the neural activity in each brain into a shared, stimulus-driven feature space significantly enhances encoding performance. This shared space isolates stimulus-driven latent features in neural activity across both subjects and electrodes, while effectively filtering out subject-specific idiosyncrasies (Haxby et al., 2011; Chen et al., 2015; Haxby et al., 2020). Our results illustrate that this shared space exhibits stronger alignment with LLM embeddings than a control model using PCA (with matching dimensionality) to aggregate signals across subjects.

SRM and other hyperalignment methods were developed, initially with fMRI, to estimate a shared information space aligned across subjects (Haxby et al., 2011; Chen et al., 2015; Guntupalli et al., 2016; Feilong et al., 2018; Van Uden et al., 2018; Haxby et al., 2020; Nastase et al., 2020; Feilong et al., 2023). ECoG acquisition presents a more challenging correspondence problem due to varying electrode numbers and placement across subjects (Owen et al., 2020). Electrode placement is often arbitrary, based on clinical considerations, yielding both redundancies and gaps in coverage, which can hamper model generalization. In most ECoG research (e.g., Goldstein et al., 2022; Cai et al., 2023; Mischler et al., 2024), electrodes are simply pooled across subjects to construct a “supersubject”. No mapping from one subject to another is attempted, and, critically, whether encoding models actually generalize across subjects has not been investigated. In the current manuscript, we extend SRM to ECoG data and demonstrate several ways in which aggregating electrode signals into a shared information space can improve encoding model performance.

The shared features estimated by SRM are linear combinations of signals across electrodes and subjects (Chen et al., 2015). To more easily interpret these signals, we reconstructed electrode-space activity from the reduced-dimension shared space. This allows us to “denoise” individual data via the shared space. We found that SRM-reconstruction improves encoding performance in most electrodes (a mean 38% improvement), particularly in brain areas associated with language processing, such as the IFG and STG. These areas both (a) contain more electrodes than other areas and (b) may be most closely entrained to linguistic features of the shared stimulus (e.g. Goldstein et al., 2022; 2024).

The vast majority of prior work fitting electrode-wise linguistic encoding models does not evaluate whether models generalize across individual subjects (Goldstein et al. 2022; Goldstein, Wang, et al. 2023; Mischler et al. 2024; cf. Zada et al. 2023). Our findings show that, by estimating both SRM and encoding models on a subset of training subjects and stimuli, the shared space can be used to build encoding models that robustly generalize to new subjects and stimuli. We show that cross-subject encoding performance via the shared space matches or even exceeds within-subject encoding performance. This generalization likely hinges on using a rich, naturalistic stimulus (like a spoken story) to obtain a diverse sampling of brain states, which ultimately yields a more robust, generalizable shared space (Haxby et al., 2011; 2020). This kind of generalization—allowing us to precisely predict neural activity in previously unseen subjects—can provide a way to circumvent the scarcity of individual-subject data, which is particularly egregious with ECoG recordings in epilepsy patients. Given a shared, naturalistic stimulus, SRM allows us to leverage previously collected data from a larger group of subjects in a single individual’s idiosyncratic electrode space—which may accelerate research on individualized brain decoding and brain-computer interfaces (e.g. Metzger et al., 2023; Willett et al., 2023).

We show that SRM improves model-based encoding performance and provides a basis for robustly generalizing encoding models across individual subjects. SRM is a data-driven, unsupervised algorithm that isolates stimulus-driven features of neural activity and aligns them across individuals. What are these stimulus-driven features? For a naturalistic, spoken language stimulus, we hypothesize that these features likely capture the structure of real-world language supporting comprehension, production, and ultimately communication. Using contextual embeddings derived from an LLM, we confirm this hypothesis by showing that SRM improves model-based encoding performance. That is, we show that by aligning neural signals across subjects, we more closely converge on the shared set of linguistic features encoded by individual brains and LLMs.

## Supplementary Information

**Table S1:**
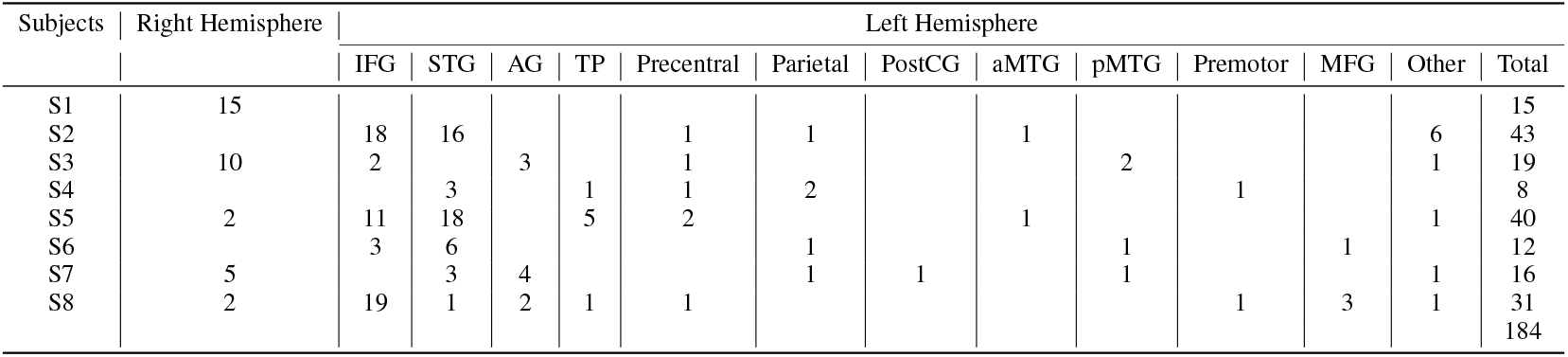
Electrode localization to different brain areas for each subject. STG: superior temporal gyrus, aMTG: anterior middle temporal gyrus, pMTG: posterior middle temporal gyrus, TP: temporal pole, AG: angular gyrus, IFG: inferior frontal gyrus, MFG: middle frontal gyrus, PostCG: postcentral gyrus.

| Subjects | Right Hemisphere | Left Hemisphere |  |  |  |  |  |  |  |  |  |  |  |  |
| --- | --- | --- | --- | --- | --- | --- | --- | --- | --- | --- | --- | --- | --- | --- |
|  |  | IFG | STG | AG | TP | Precentral | Parietal | PostCG | aMTG | pMTG | Premotor | MFG | Other | Total |
| S1 | 15 |  |  |  |  |  |  |  |  |  |  |  |  | 15 |
| S2 |  | 18 | 16 |  |  | 1 | 1 |  | 1 |  |  |  | 6 | 43 |
| S3 | 10 | 2 |  | 3 |  | 1 |  |  |  | 2 |  |  | 1 | 19 |
| S4 |  |  | 3 |  | 1 | 1 | 2 |  |  |  | 1 |  |  | 8 |
| S5 | 2 | 11 | 18 |  | 5 | 2 |  |  | 1 |  |  |  | 1 | 40 |
| S6 |  | 3 | 6 |  |  |  | 1 |  |  | 1 |  | 1 |  | 12 |
| S7 | 5 |  | 3 | 4 |  |  | 1 | 1 |  | 1 |  |  | 1 | 16 |
| S8 | 2 | 19 | 1 | 2 | 1 | 1 |  |  |  |  | 1 | 3 | 1 | 31 |
|  |  |  |  |  |  |  |  |  |  |  |  |  |  | 184 |

**Figure S1:**
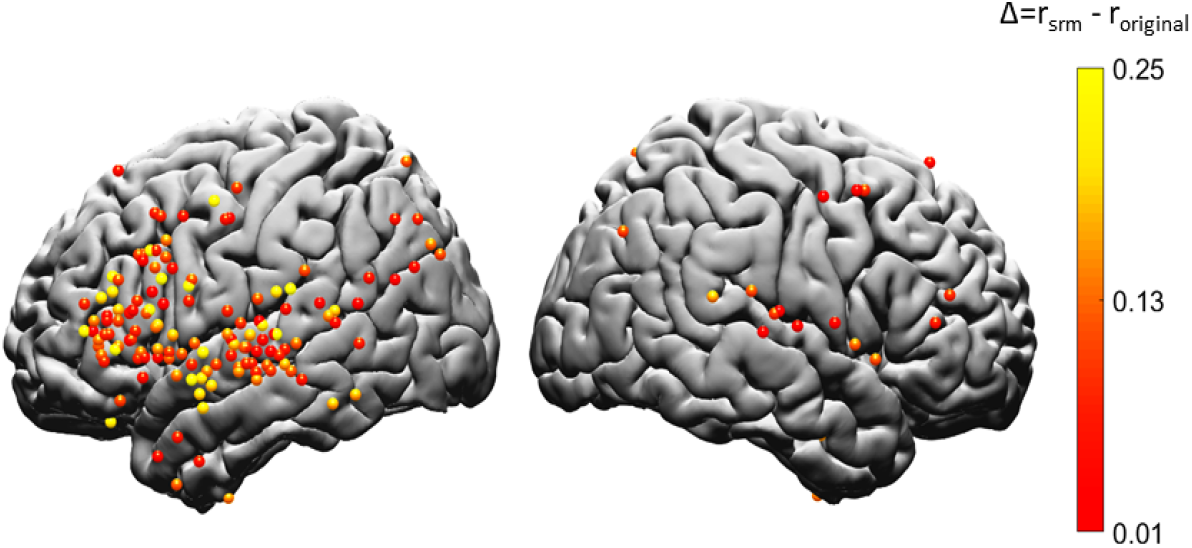
Electrode-wise differences in encoding model performance between SRM-reconstructed data and the original electrode data. Differences were computed between the respective maximum encoding performance values across lags for SRM-reconstructed and original electrode data.

**Figure S2:**
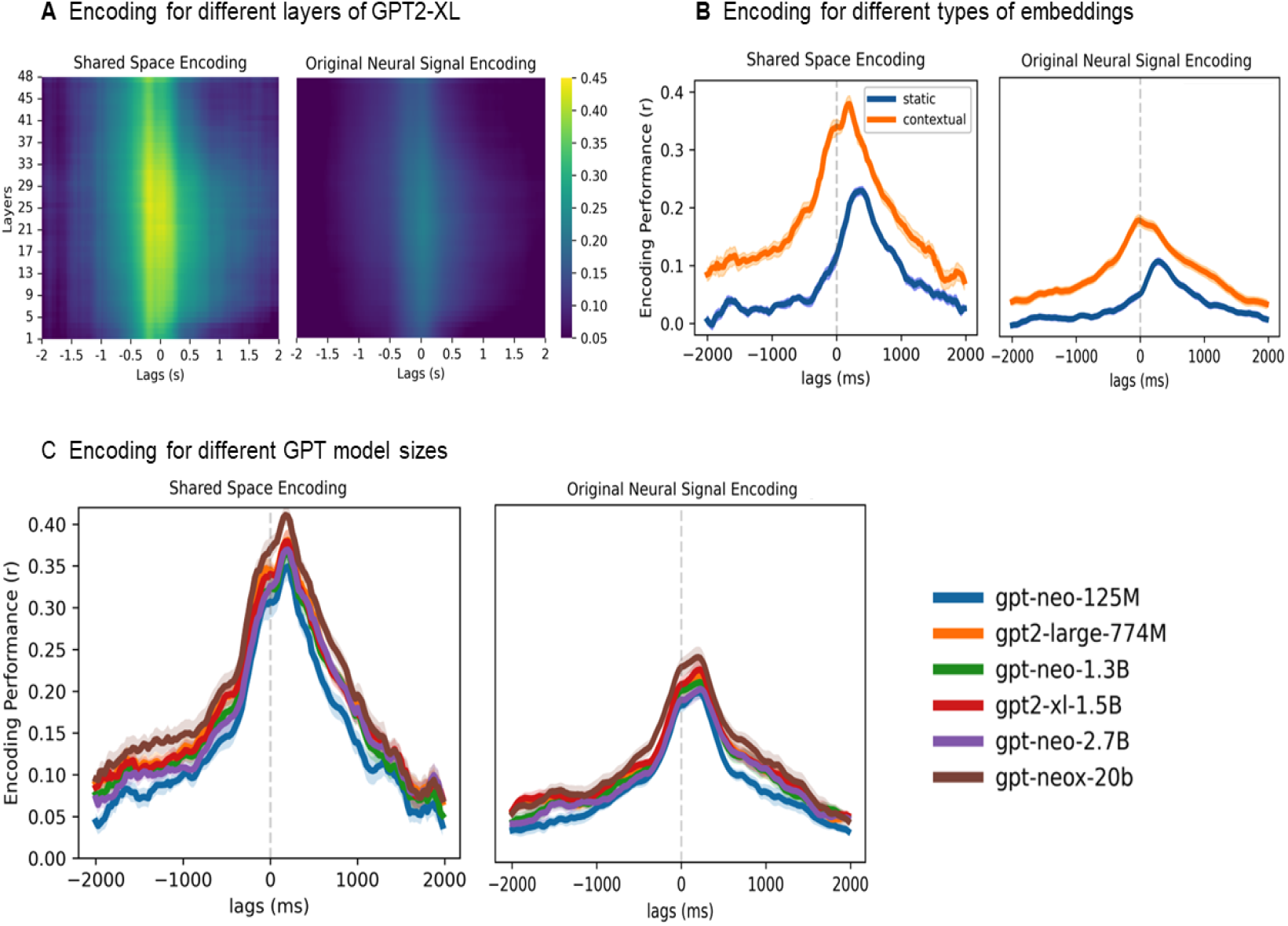
Exploring SRM encoding across different model features. (**A**) Encoding performance across all the layers of GPT-2 XL for both shared space (left) and original electrode data (right). (**B**) Comparison of encoding performance for contextual embeddings from GPT-2 XL (orange) and non-contextual embeddings from GloVe (blue) in both the shared space (left) and original electrode data (right). (**D**) Encoding performance across different sizes of GPT models for both shared space and original electrode data. In all cases, the error bands indicate the standard error of the mean across folds.

## Code availability

https://github.com/pritamarnab/SRM-Encoding

